# De novo protein design by deep network hallucination

**DOI:** 10.1101/2020.07.22.211482

**Authors:** Ivan Anishchenko, Tamuka M. Chidyausiku, Sergey Ovchinnikov, Samuel J. Pellock, David Baker

## Abstract

There has been considerable recent progress in protein structure prediction using deep neural networks to infer distance constraints from amino acid residue co-evolution^1–3^. We investigated whether the information captured by such networks is sufficiently rich to generate new folded proteins with sequences unrelated to those of the naturally occuring proteins used in training the models. We generated random amino acid sequences, and input them into the trRosetta structure prediction network to predict starting distance maps, which as expected are quite featureless. We then carried out Monte Carlo sampling in amino acid sequence space, optimizing the contrast (KL-divergence) between the distance distributions predicted by the network and the background distribution. Optimization from different random starting points resulted in a wide range of proteins with diverse sequences and all alpha, all beta sheet, and mixed alpha-beta structures. We obtained synthetic genes encoding 129 of these network hallucinated sequences, expressed and purified the proteins in E coli, and found that 27 folded to monomeric stable structures with circular dichroism spectra consistent with the hallucinated structures. Thus deep networks trained to predict native protein structures from their sequences can be inverted to design new proteins, and such networks and methods should contribute, alongside traditional physically based models, to the de novo design of proteins with new functions.

## Introduction

Deep learning methods have shown considerable promise in protein engineering. Networks with architectures borrowed from language models have been trained on amino acid sequences, and been used to generate new sequences without considering protein structure explicitly ^4,5^. Other methods have been developed to generate protein backbones without consideration of sequence ^6^, and to identify amino acid sequences which either fit well onto specified backbone structures ^7–9^ or are conditioned on low-dimensional fold representation ^10^; models tailored to generate sequences and/or structures for specific protein families have also been developed ^11–14^. However, none of the models described to date address the classical de novo protein design problem: generating new sequences predicted to fold to new structures.

Deep neural networks trained to predict distances between amino acid residues in protein 3D structures from amino acid sequence information have increased the accuracy of protein structure prediction ^1–3^. These models take as input large sets of aligned sequences, and a major contributor to distance prediction accuracy is the extent of co-evolution between the amino acid identities at pairs of positions: two positions where the amino acid identity changes in the alignment are correlated are likely to be in contact in the protein 3D structure. Following up on an initial observation by AlphaFold in CASP13 ^15^, we found that the trRosetta deep neural network trained using multiple sequence information could consistently predict structure quite accurately for de novo designed proteins from just a single sequence--ie, in the complete absence of co-evolution information ^3^. The trRosetta model also predicted the effects of amino acid substitutions on folding consistent with biophysical expectation. These results suggested that during training the network was learning fundamental relationships between protein sequence and structure.

We wondered if the information stored in the many parameters of the trained networks could be used to design new protein structures with new sequences. Methods such as Google’s DeepDream ^16^ take networks trained to recognize faces and other patterns in images, and invert these by taking arbitrary input images and adjusting them to be more strongly recognized as faces (or other patterns) by the network--the resulting images are often referred to as hallucinations because they do not represent any actual face, but what the neural network views as an ideal face. We decided to take a similar approach to explore whether networks trained to predict structures from sequences could be inverted to generate brand new “ideal” protein sequences and structures.

We started from the trRosetta deep neural network which predicts distributions of distances and orientations between all pairs of residues from a set of aligned protein sequences from a protein family (Fig 1A); in benchmark tests this network outperformed other methods ^3^. Instead of inputting a naturally occurring sequence, we instead generated completely random 100 amino acid sequences, and fed these to the network (Fig 1B). As expected for random sequences, which have a vanishingly small probability of folding to a defined structure, the distance distributions were diffuse and much less featured than those obtained with actual protein sequences. We then sought to optimize the sequences such that the network predicted distance map was as different as possible (had the highest Kullback-Leibler divergence) from the background distribution (Fig 1B,C and methods). For each sequence, we carried out a Monte Carlo simulated annealing trajectory in sequence space: each step consists of introducing a random mutation (single amino acid substitution) somewhere in the sequence, predicting the distance map of the mutated sequence using the network, and accepting the move based on the change in the KL-divergence to the background distribution according to the standard Metropolis criterion (Fig 1C and methods). The increase in KL-divergence aggregated over all 2,000 simulation trajectories is shown in Fig 1D; in almost all cases after ~20,000 Monte Carlo steps the resulting distance maps were at least as featured (non-uniform) as those predicted for native sequences and structurally confirmed de novo proteins designed using Rosetta. The predicted distance maps become progressively sharper during the course of the simulations, and trajectories started from different random sequences resulted in very different sequences and structures (Fig 1E). We converted the final sharpened distance maps into protein 3D structures by direct minimization with trRosetta^3^.

**Figure 1:**
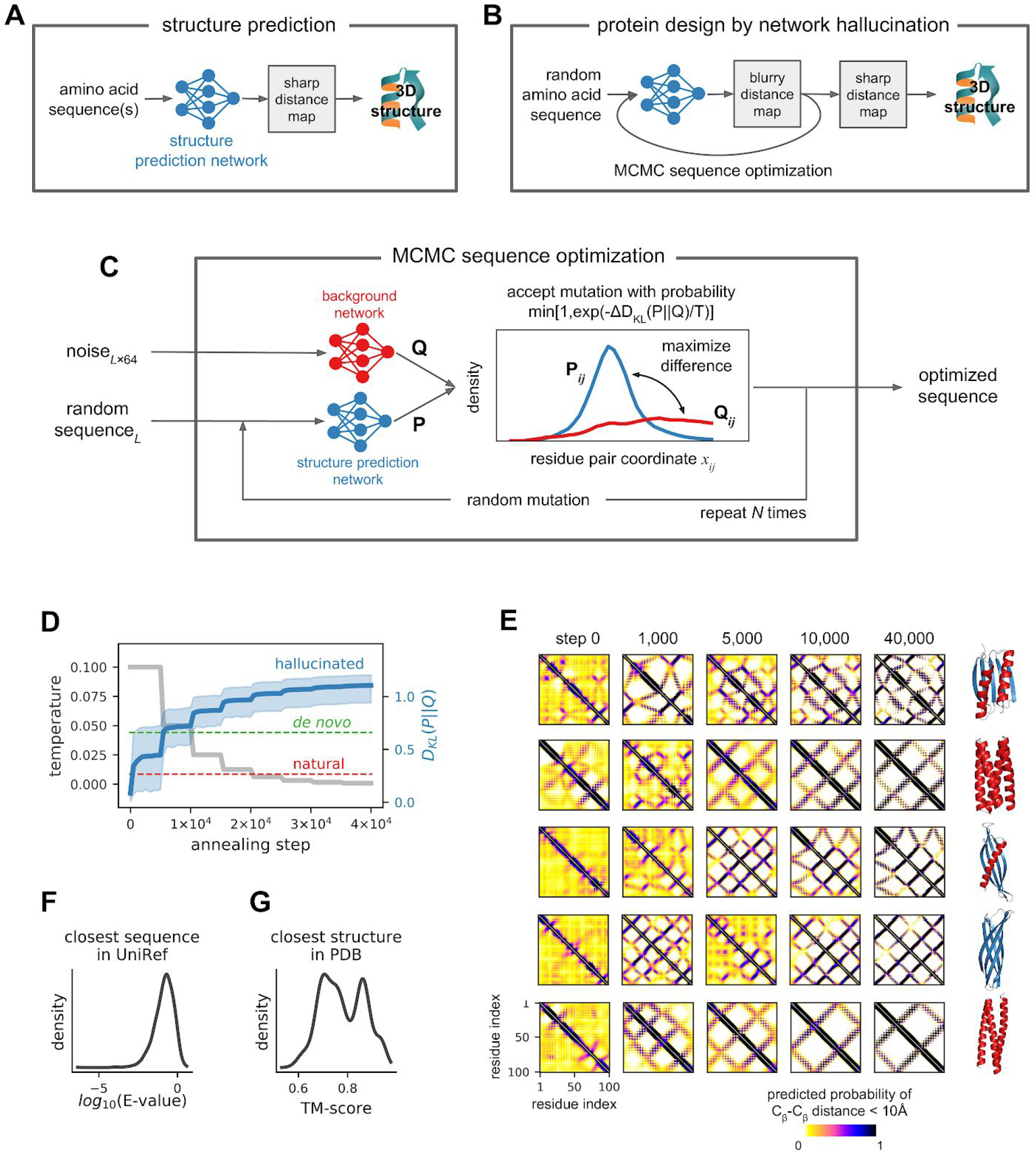
Overview of protein hallucination approach. **A)** In structure prediction using trRosetta and other recent methods, a deep neural network is used to predict inter-residue geometries (reliable predictions have sharp 2D distance and orientation maps) from a single sequence or a multiple sequence alignment, and then the 3D structure is reconstructed by constrained minimization. **B)** Network predictions for a random sequence are not confident (blurry 2D maps); to transform a random sequence into one encoding a new folded protein, we introduce multiple single amino acid substitutions into the sequence using Markov chain Monte Carlo algorithm, optimizing the sharpness of the 2D maps. **C)** Schematic of the MCMC procedure. **D)** Annealing trajectories averaged over 2,000 runs show a monotonic increase in the KL-divergence (contrast of the distance maps) with increasing Monte Carlo optimization. The mean and 0.01,0.99 quantiles are shown in magenta; temperature profile is shown in grey. **E)** Distance maps become progressively sharper along the Monte Carlo trajectories as exemplified by five hallucinated sequences with different protein structure topologies. **F)** Hallucinated sequences are unrelated to the naturally occurring protein sequences in the UniRef90 database: median BLAST E-value of the closest hit is 0.17. **G)** Hallucinated structures range in similarity to the protein structures in the PDB with average TM-scores to the closest match of 0.78.

We used this approach to generate two thousand new proteins with sequences predicted by the trRosetta network to fold into well defined structures, and compared their sequences and structures to native protein sequences and structures. The similarity of the hallucinated sequences to native protein sequences was very low, with only very short alignable regions and best Blast e-values to the Uniprot database of ~0.1 (Fig 1F). Just as simulated images of cats generated by deep network hallucination are clearly recognizable as cats, but differ from the images the network was trained on, the predicted structures resemble but are not identical to native structures in the PDB, with TM-align scores of 0.6-0.9 (Fig 1G).

The hallucinated sequences and their associated structures are quite diverse--different Monte Carlo trajectories converge on different sequence-structure pairs (Fig 2A and B). A 2D map of the space spanned by the structures (Fig 2B) was generated by multidimensional scaling of their pairwise 3D structural similarity (TM-score, see Methods). The structures span all alpha, all beta and mixed alpha-beta fold classes, and within these classes a total of 27 different sub-folds were identified at a TM-align clustering threshold of 0.8; representative examples are shown in Fig 2C.

**Figure 2:**
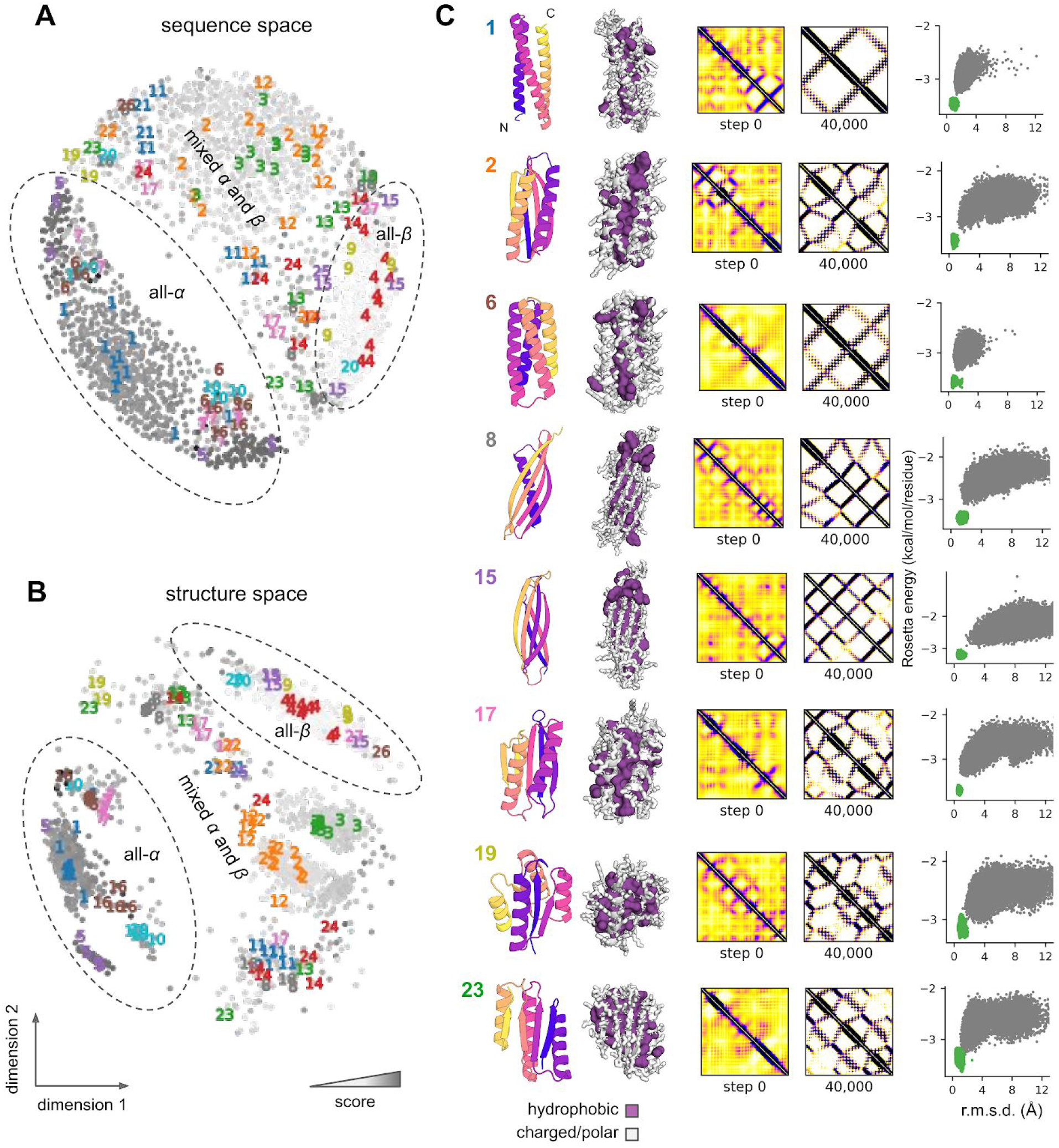
Overview of computational results. **A)** Multidimensional scaling plot of the sequence space covered by the 2,000 hallucinated proteins; BLAST bit-score was used to measure the distance between proteins. Each grey dot represents one design color-coded by the score from the network (darker grey color corresponds to higher score). 129 experimentally tested designs belong to 27 structural clusters shown by colored numbers. **B)** Multidimensional scaling plot of the structural space covered by the 2,000 hallucinated proteins; (1 – TM-score) was used to measure the distance between proteins. Each grey dot represents one design; the gray-scale indicates the score from the network (darker grey corresponds to higher score). The 129 experimentally tested designs fall into 27 structural clusters shown by colored numbers. **C)** Examples of hallucinated designs of various topologies. First column, ribbon depiction of protein backbone colored from blue (N-terminus) to red (C-terminus); second column, hydrophobic core; third column, distance maps at the beginning and end of hallucination trajectory, and fourth column, folding energy landscapes from large scale Rosetta *ab initio* structure prediction calculations; points represent lowest-energy structures sampled starting from an extended chain (green points) and starting from the hallucinated design model (grey points). The energy landscapes funnel into the energy minimum corresponding to the designed structure, providing independent, albeit in silico, evidence that the hallucinated sequences encode the hallucinated structures.

A striking feature of the hallucinated structures is that they resemble the “ideal” proteins generated by de novo protein design more than native proteins, despite the fact that the network was trained on the latter. De novo protein design research over the past 10 years has sought to distill the key features of protein structures and protein sequence-structure relationships using physically based models like Rosetta, and then has used these models to design idealized structures that embody these features based on the principle that proteins fold to their lowest free energy states ^17,18^. Both de novo designed proteins and the hallucinated proteins generated here have regular alpha helices and beta sheets, and lack the long loops and other idiosyncrasies of native protein structures. Remarkably, in training the deep neural network on large numbers of (irregular) native protein structures, it learned to encode ideal protein structure properties very similar to expert protein designers using more traditional scientific approaches. Of course, the representation of this knowledge is very different; in the case of traditional de novo protein design, the knowledge is manifested in the energy function and sampling methods in protein design software such as Rosetta, whereas in the deep network it is distributed amongst the millions of network parameters (in a much less interpretable fashion).

We next set out to determine whether the hallucinated sequences actually adopt folded structures. We first used in silico protein folding simulations using Rosetta^19^ to assess the extent to which the sequences encode the structures according to the Rosetta forcefield ^20^. This is a completely orthogonal test as the network was trained exclusively on native protein structures, and has no access to the Rosetta energy function. We generated folding energy landscapes using large scale de novo folding simulations starting from an extended chain for 129 of the hallucinated proteins spanning a wide range of sequences and structures. For 82 out of 129 of these, the lowest energy structures found in the simulations were close to the corresponding hallucinated structures with Ca-RMSD less than 3.0 A, and for all 129, the lowest energy structure sampled starting from the design model was lower than in energy than any obtained starting from an extended chain. Thus according to the Rosetta physically based energy model, the network generated sequences do indeed encode the corresponding structures.

We next sought to determine whether the hallucinated proteins actually fold in the real world by obtaining synthetic genes for the 129 proteins, and expressing and purifying them from E coli (see Methods). All 129 of the proteins were solubly expressed. 32 had clear peaks in size exclusion chromatography with the apparent molecular weights of the monomer, and 3 had clear peaks corresponding to small oligomers (Sup fig 2). For 27 of these, the peaks corresponding to the monomeric or small oligomeric species were examined by circular dichroism (CD) spectroscopy. In all cases, the CD spectra were consistent with the target structures (Figs 3 and 4, E), with the characteristic profiles of all alpha helical proteins for the all alpha helical designs (Fig 3), and of alpha-beta proteins for the alpha-beta designs (Fig 4). Most of the designs were thermostable, with apparent melting temperatures above 70°C (Figs 3 and 4, F). The alpha-beta designs in figure 4 are particularly stable; none undergo unfolding transitions up to 95°C. Taken together, these data indicate that many of the network hallucinated proteins fold up into stable monomeric structures with the predicted secondary structure content.

**Figure 3.**
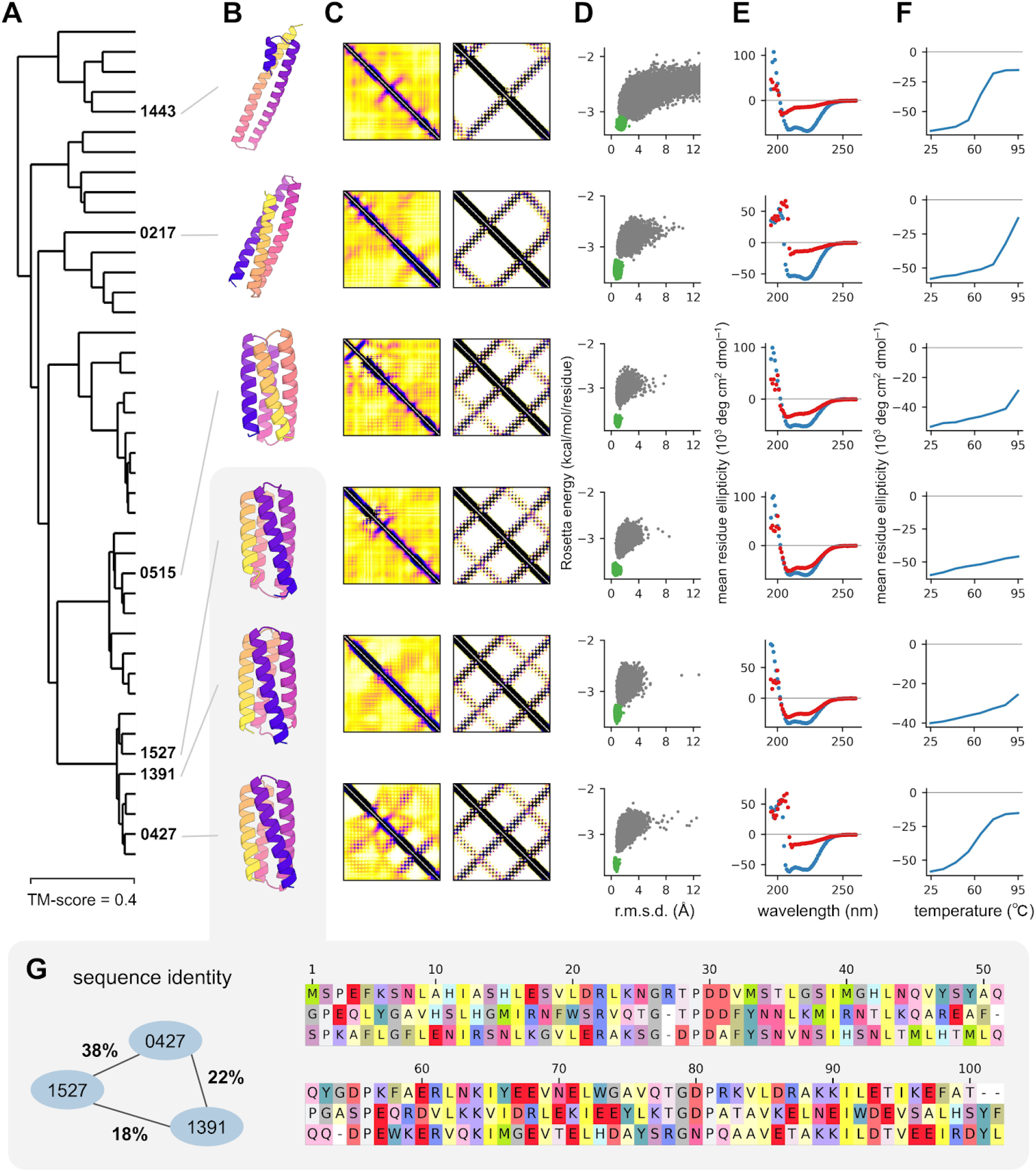
Experimental characterization of alpha-helical hallucinations. **A)** Dendrogram showing 42 all-alpha designs clustered by structural similarity (TM-score); thermostable designs with CD spectra consistent with the target structure are labeled by their IDs. **B)** 3D structure models of the hallucinated designs. **C)** Distance maps before and after the hallucination process. **D)** *ab initio* folding funnels from Rosetta. **E)** Circular dichroism spectra at 25 (blue) and 95°C (red). **F)** Temperature dependence of circular dichroism signal at 220nm between 25°C and 95°C. **G)** Albeit structurally similar (Ca-RMSD ~ 1A), designs 0427, 1391 and 1527 have very different amino acid sequences with pairwise sequence identities not exceeding 38%.

**Figure 4.**
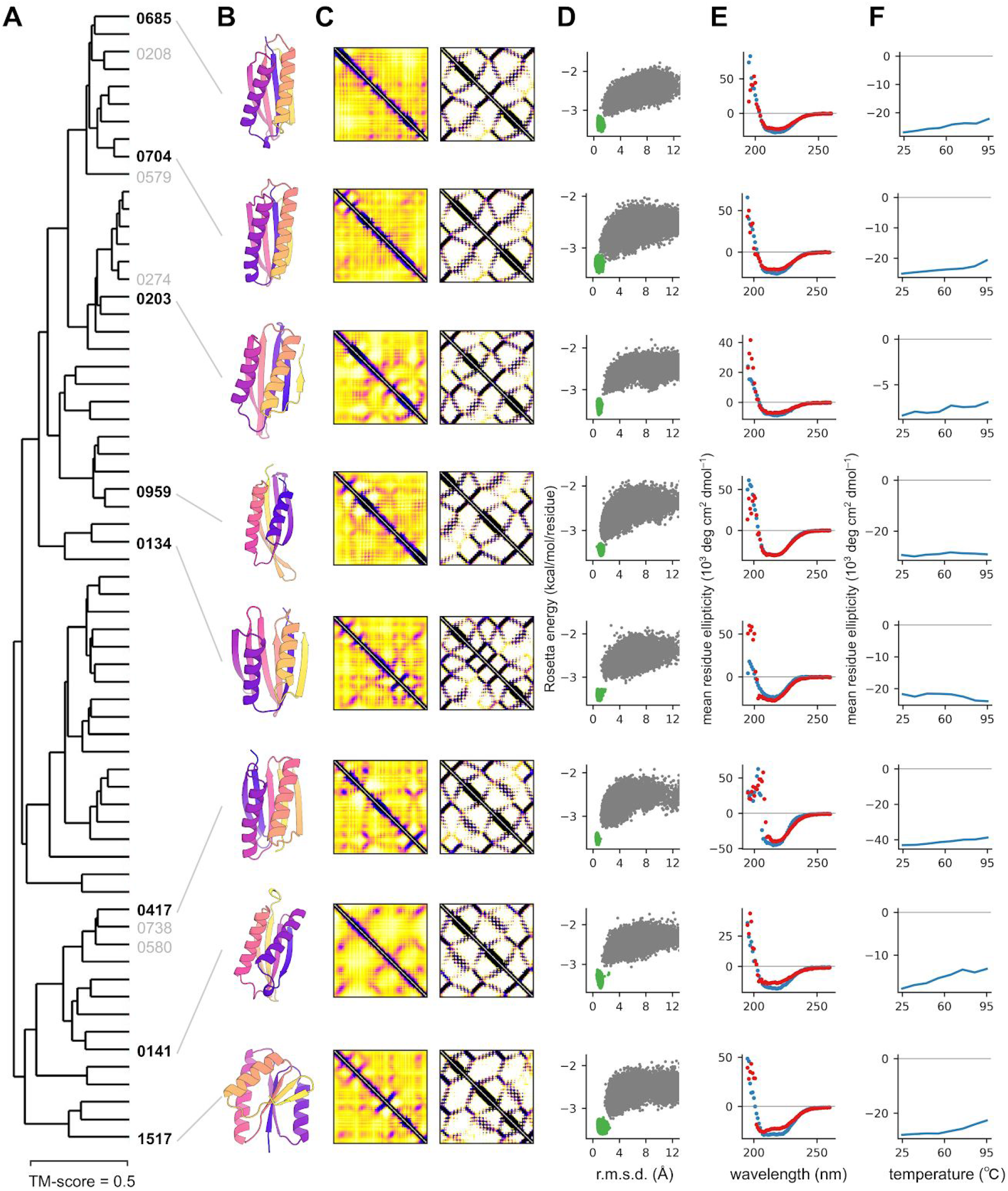
Experimental characterization of mixed alpha and beta hallucinations. Columns are labeled as in Figure 3. Additional examples of stable alpha and beta designs marked in grey in panel **A** are shown in Fig S3.

The designs that adopt monomeric structures with the expected CD spectra fall into 11 of the 27 structure classes noted above. Representative examples of all alpha and alpha-beta experimentally validated designs are shown in Figs 3 and 4, respectively. In each class, the designs span a wide range of topologies (dendrograms on left). The distance predictions for the random starting sequences (panels C, left side) have few features and do not differ substantially between structure classes, but are clearly distinct following hallucination optimization (panels C, right side). All of the sequences are predicted by Rosetta large scale energy calculations (panels D) to have funneled landscapes leading into the target structure, but detailed confirmation of this will await experimental high resolution structure determination.

## Conclusion

Our results demonstrate that a deep neural network trained exclusively on native sequences and structures can generalize to create new proteins, with sequences unrelated to those of native proteins, that fold into stable monomeric structures. The resemblance--in the regularity of the secondary structures, shortness of the loops, etc-- to idealized proteins designed, using programs such as Rosetta, based on the principle that proteins fold to their lowest energy structures is remarkable--both approaches appear to have extracted the same core features, albeit representing them within the corresponding models in very different ways (in the millions of parameters in the network in the first case, and in the very much smaller number of parameters of the force field, in the second). The hallucinated proteins are monomeric, stable, have the expected secondary structure, and are strongly predicted to fold to the target structure by Rosetta in completely orthogonal calculations (we did not use Rosetta in any way for either sequence generation or selection for experimental characterization), suggesting that they fold close to the target structure. We are attempting to solve the structures of a subset of these proteins by x-ray crystallography and NMR.

Our work opens up a large set of exciting avenues to explore. On the sampling side, the Monte Carlo approach could perhaps be made more efficient by direct gradient based minimization by tracing the gradients back to the inputs, as in image hallucination work. The loss function can be generalized to include specific structural features, for example binding motifs or catalytic sites, around which the network can hallucinate new protein inhibitors or enzyme catalysts. Unlike traditional protein design calculations, where properties of the target scaffold such as the overall topology and/or the secondary structure element lengths and locations are specified in advance, through a structure “blueprint” or other approach, the ability of the network to hallucinate plausible protein structures from scratch makes building a supporting scaffold around a desired functional site much more straightforward since the structure need not be mapped out in advance; the network can also come up with a wide range of different protein topology solutions for a given problem. More generally, our work demonstrates the power of generative deep learning approaches for molecular design, which will undoubtedly continue to grow over the coming years.

## Methods

### Approach

The general protein design problem can be formulated in probabilistic terms as the finding of mutually compatible sequence-structure pairs such that the joint probability P(sequence, structure) is maximized. Using the chain rule for probabilities:

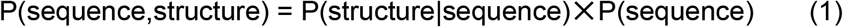

The first term on the right, P(structure|sequence), is related to the protein structure prediction problem where one seeks for the most probable structure for a given protein sequence. The second term P(sequence) accounts for all possible intrinsic biases in amino acid composition not related to structure. We use a pre-trained neural network previously developed for protein structure prediction to approximate the P(structure|sequence) term, and then maximize the joint probability P(sequence, structure) in sequence space by simulated annealing.

### Networks & objective function

The *trRosetta* protein structure prediction network, described in detail elsewhere^3^, is a 2D residual-convolutional neural network which takes 1- and 2-site features derived from a multiple sequence alignment or a single sequence as an input and produces a 2D output describing distances and orientations for all residue pairs in a protein in a probabilistic manner: for every residue pair (i,j), these generated maps contain predicted probability distributions over the Cβ-Cβ distance and 5 inter-residue angles (comprising the full set of 6 rigid-body degrees of freedom). When accurate, such 2D predictions can be straightforwardly translated into a 3D structure by direct minimization ^2,3^.

Random sequences give diffuse predictions, while existing de novo designs produce peaked distributions with low variance^3^. To quantify the quality of network predictions and as the optimization objective we use the information gain (the Kullback-Leibler divergence *D_KL_*) in the predicted structure distributions relative to the background distributions P(structure) when the sequence is not known:

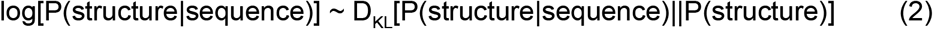

To approximate P(structure) we trained a separate *background* network which is similar in architecture to *trRosetta* but the inputs to the network are sequence-agnostic and are represented by random gaussian noise of shape *L*×64. Such a network effectively generates inter-residue probability distributions which carry the information on all protein structures in the PDB and can loosely be viewed as a generic “molten globule” state.

### Amino acid composition bias

The P(sequence) term in Eq. (1) is responsible for keeping the amino acid composition of the hallucinated designs within the viable region of the protein sequence space. This was achieved by calculating frequencies of each of the 20 standard amino acids *f_a_* within the generated sequences and comparing them against corresponding frequencies 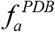 collected over high-res X-ray structures from the Protein Data Bank. To keep *f_a_* close to 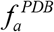, negative KL-divergence was used:

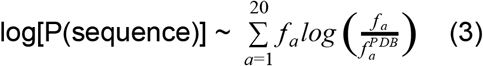

During sequence generation, we also disallow cysteines to avoid the experimental complications of disulfide bonds by sampling from 19 of the remaining standard amino acids only.

### Sequence optimization

Given Eqs.(2-3), maximizing the joint probability of sequence and structure in Eq. (1) is equivalent to maximizing the following objective:

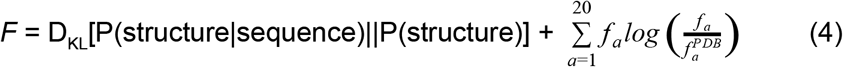

P(structure) is a fixed distribution which depends only on the protein length and is generated only once at the beginning of simulations; 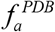 is fixed too; hence the optimization focuses on maximizing D_KL_. The design procedure starts off with picking a random amino acid sequence of a given length *L* (*L* = 100 throughout the study), passing it through trRosetta and background networks and calculating the objective *F* according to Eq.(4). Next, random single amino acid mutations are introduced into the sequence -- one at a time -- and each mutated sequence is again passed through the trRosetta network. The decision whether the mutation at step *i* is accepted or not is based on the Metropolis criterion. We calculate the acceptance probability *A_i_* of the candidate sequence by comparing objectives (Eq.4) at steps *i* and *i*−1

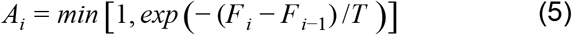

and accept the mutation only if *A_i_* is smaller than a uniform random number *u* ∈ [0, 1]. A total of 40,000 mutations are probed to generate a single design. Temperature *T* is set to 0.1 at the beginning of the trajectory and then is reduced by half every 5,000 steps.

### Design selection

After generating 2,000 hallucinated sequences and folding them using *trRosetta* minimization protocol, the resulting structures were compared to each other in terms of the template modeling score (TM-score) ^21^. The pool of designs was then split into clusters at TM-score threshold equal to 0.8. We then picked clusters with at least 7 members and scored the designs separately within each cluster by a two-component scoring function:

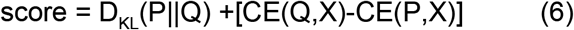

P and Q in Eq.(6) are shortcuts to P(structure|sequence) and P(structure) as predicted by *trRosetta* and *background* networks respectively, CE stands for cross-entropy, and X is the 2D representation of the realized 3D structure in terms of residue-residue distances and orientations. The first component to the *score* in Eq.(6) -- D_KL_(P‖Q) -- is the same as in Eq.(2): it measures the quality of the generated distance and orientation maps with no regard to the 3D structure. The second term [CE(Q,X)-CE(P,X)] takes into account the realized 3D structure of the hallucinated design and checks how well it actually fits into the original *trRosetta* predictions vs *background*. Top-scoring designs from each cluster were manually inspected, and finally 129 hallucinated sequences from 27 structural clusters (no more than 10 designs per cluster) were selected for experimental testing.

### Protein Expression and Purification

Genes coding for the selected 129 designs were synthesized and cloned into pET28 b(+) expression vector with an additional 21-residue N-terminal sequence containing a His-tag and thrombin cleavage site to aid purification (full sequence: MGSSHHHHHHSSGLVPRGSHM). These plasmids were purchased from Genscript and expressed in E. coli BL21(DE3) cells. Starter cultures were grown overnight at 37°C in Luria-Bertani (LB) medium with added antibiotic (50 μg/ml Kanamycin). These overnight cultures were used to inoculate either 50mL (for screening) or 500 mL (for crystallography) of Studier autoinduction media^22^ supplemented with antibiotic, and grown overnight. Cells were harvested by centrifugation and resuspended in 25 mL lysis buffer (20 mM imidazole in PBS containing protease inhibitors), and lysed by microfluidizer. PBS buffer contained 20mM NaPO4, 150mM NaCl, pH 7.4. After removal of insoluble pellets, the lysates were loaded onto nickel affinity gravity columns to purify the designed proteins by immobilized metal-affinity chromatography (IMAC).

### Size-exclusion Chromatography for Screening

To determine oligomeric state, following IMAC, all designs showed soluble expression on SDS-PAGE gels and therefore were further purified by size-exclusion chromatography on ÄKTAxpress (GE Healthcare) using a Superdex 75 10/300 GL column (GE Healthcare) in PBS buffer. The monomeric or smallest oligomeric fractions of each run (typically eluting at the 14 mL mark) were collected and immediately analyzed by circular dichroism (CD) or flash frozen in liquid N2 for later analysis.

### Circular Dichroism Experiments

To determine secondary structure and thermostability of the designs far-ultraviolet CD measurements were carried out with an JASCO 1500. 260 to 195 nm wavelength scans were measured at every 10°C interval from 25°C to 95°C. Temperature melts monitored dichroism signal at 220 nm in steps of 2°C/minute with 30s of equilibration time. Wavelength scans and temperature melts were performed using 0.35 mg/ml protein in PBS buffer (20mM NaPO4, 150mM NaCl, pH 7.4) with a 1 mm path-length cuvette. Chemical denaturation experiments with guanidinium hydrochloride (GuHCl) were performed using an automatic titrator with a protein concentration of 0.035 mg/ml and a 1 cm path-length cuvette with stir bar. The GuHCl concentration was determined by refractive index in PBS buffer. The denaturation process monitored dichroism signal at 220 nm in steps of 0.2 M GdmCl with 1 min mixing time for each step, at 25°C. Protein concentrations were determined by absorbance at 280 nm measured using a NanoDrop spectrophotometer (Thermo Scientific) using predicted extinction coefficients^23^.

## Supplementary figures

**Figure S1.**
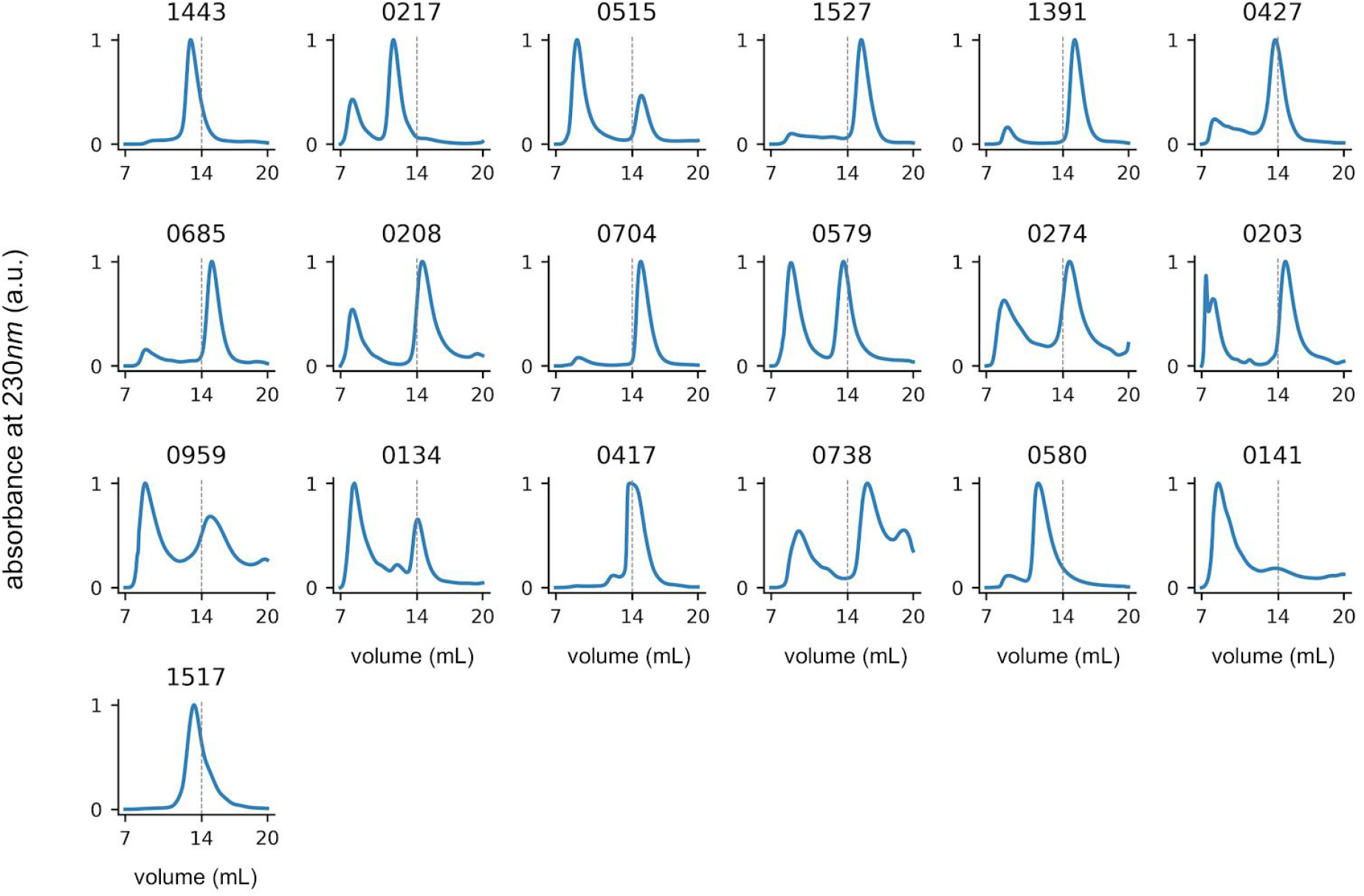
Size-exclusion chromatography traces for the designs shown in Figs 3, 4 and S2. Vertical line at 14mL shows the approximate location of the monomeric or smallest oligomeric fractions which were used for circular dichroism measurements.

**Figure S2.**
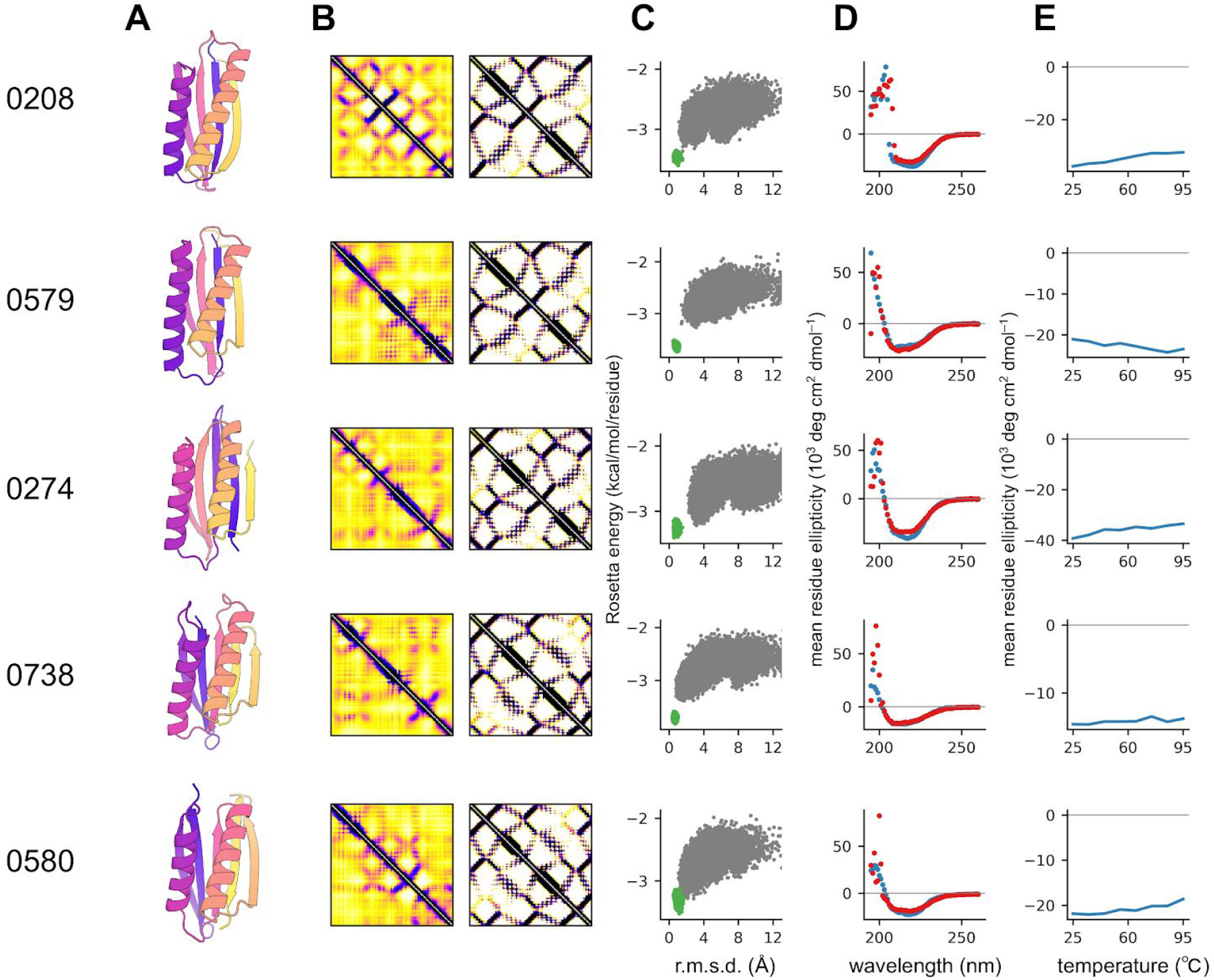
Additional examples of mixed alpha and beta hallucinations (see Fig 4 for details)

